# Petascale neural circuit reconstruction: automated methods

**DOI:** 10.1101/2021.08.04.455162

**Authors:** Thomas Macrina, Kisuk Lee, Ran Lu, Nicholas L. Turner, Jingpeng Wu, Sergiy Popovych, William Silversmith, Nico Kemnitz, J. Alexander Bae, Manuel A. Castro, Sven Dorkenwald, Akhilesh Halageri, Zhen Jia, Chris Jordan, Kai Li, Eric Mitchell, Shanka Subhra Mondal, Shang Mu, Barak Nehoran, William Wong, Szi-chieh Yu, Agnes L. Bodor, Derrick Brittain, JoAnn Buchanan, Daniel J. Bumbarger, Erick Cobos, Forrest Collman, Leila Elabbady, Paul G. Fahey, Emmanouil Froudarakis, Daniel Kapner, Sam Kinn, Gayathri Mahalingam, Stelios Papadopoulos, Saumil Patel, Casey M. Schneider-Mizell, Fabian H. Sinz, Marc Takeno, Russel Torres, Wenjing Yin, Xaq Pitkow, Jacob Reimer, Andreas S. Tolias, R. Clay Reid, Nuno Maçarico da Costa, H. Sebastian Seung

## Abstract

3D electron microscopy (EM) has been successful at mapping invertebrate nervous systems, but the approach has been limited to small chunks of mammalian brains. To scale up to larger volumes, we have built a computational pipeline for processing petascale image datasets acquired by serial section EM, a popular form of 3D EM. The pipeline employs convolutional nets to compute the nonsmooth transformations required to align images of serial sections containing numerous cracks and folds, detect neuronal boundaries, label voxels as axon, dendrite, soma, and other semantic categories, and detect synapses and assign them to presynaptic and postsynaptic segments. The output of neuronal boundary detection is segmented by mean affinity agglomeration with semantic and size constraints. Pipeline operations are implemented by leveraging distributed and cloud computing. Intermediate results of the pipeline are held in cloud storage, and can be effortlessly viewed as images, which aids debugging. We applied the pipeline to create an automated reconstruction of an EM image volume spanning four visual cortical areas of a mouse brain. Code for the pipeline is publicly available, as is the reconstructed volume.

## Introduction

Progress in 3D electron microscopy (EM) can be seen in the increasing size of imaged volumes from the *Drosophila* brain. A reconstruction of seven medulla columns was based on 0.13 TB of raw image data (Takemura et al. 2017). A “hemibrain” reconstruction was based on 20 TB of raw image data (Scheffer et al. 2020). A 100 TB dataset of an entire *Drosophila* brain (Zheng et al. 2018) has been automatically reconstructed and is currently being proofread (P. H. Li et al. 2019; S. Dorkenwald et al. 2020).

Scaling up to even larger EM volumes is of clear interest for tackling mammalian brains. As a step in this direction, we have released a new automated reconstruction of a near cubic millimeter of visual cortex from a P87 male mouse (MICrONS Consortium et al. 2021). The reconstruction is based on roughly 1.5 petapixels of raw images acquired by serial section EM (Yin et al. 2020), a popular method of 3D EM. This is comparable in size to a recently released dataset acquired from human temporal cortex (A. Shapson-Coe et al. 2021). Both petascale datasets are much larger than recent cortical reconstructions, which were based on 0.15 TB (Motta et al. 2019) and 4.9 TB (N. L. Turner et al. 2020; S. Dorkenwald et al. 2019; C. M. Schneider-Mizell et al. 2020) datasets. The present dataset is accompanied by the visual responses of 75,000 cells in the volume recorded *in vivo* by calcium imaging prior to electron microscopy.

In the plane tangential to the cortical sheet, our mouse cortical volume spans roughly 1.3 mm (mediolateral) by 0.88 mm (anterior-posterior), including portions of primary visual cortex (V1) and higher visual areas LM, RL, and AL (Wang and Burkhalter 2007). We will also refer to these cortical areas as VISp, VISlm, VISrl, and VISal (Swanson 2018). In the radial direction, normal to the cortical sheet, the volume spans roughly 0.8 mm. These dimensions were estimated by registering the EM image to a light microscopic image acquired from the same volume *in vivo*. The volume includes all cortical layers, though extremes of L1 and L6 were omitted, where the alignment of the serial section EM images could not handle the increased frequency of imaging artifacts.

The goal of this paper is to describe the computational pipeline that was used to process the petascale dataset. Serial section images were aligned into a 3D image stack using a new approach based on convolutional nets that were trained to generate the nonsmooth transformations required to align images of sections that contain cracks and folds (Mitchell et al. 2019). Neuronal boundaries were detected by a convolutional net trained with extensive data augmentation for improving robustness to misalignments and image defects (Lee et al. 2017; S. Dorkenwald et al. 2019), as well as ground truth that was curated to cover a diverse set of cellular structures. A convolutional net was applied to classify voxels into semantic categories such as axon, dendrite, and soma. The output of neuronal boundary detection was segmented by watershed (Zlateski and Seung 2015) and mean affinity agglomeration (Lee et al. 2017; Funke et al. 2019; Lu, Zlateski, and Seung 2021) with semantic and size constraints. Convolutional nets were used to detect synaptic clefts, and assign presynaptic and postsynaptic partners to each cleft (Nicholas L. Turner et al. 2020). Pipeline operations were implemented using several frameworks that we developed for applying distributed and cloud computing to large 3D images (Nicholas L. Turner et al. 2020; J. Wu et al. 2021; Lu, Zlateski, and Seung 2021). Pipeline operations communicated with each other by using cloud storage to hold intermediate results, which could be viewed effortlessly for debugging purposes.

The aligned images and automated reconstruction have been publicly released at the MICrONS Explorer (microns-explorer.org) site (MICrONS Consortium et al. 2021). Code is also publicly available (see Data and Code Availability). While the automated reconstruction contains errors, these can be interactively corrected by human experts to yield accurate semiautomated reconstructions of neurons. This process is called “proofreading,” and is described in a companion paper.

## Image acquisition and alignment

Images were acquired using techniques described elsewhere (Yin et al. 2020; MICrONS Consortium et al. 2021). It took 12 days to cut the brain volume into over 27,000 sections, which were collected onto GridTape (Phelps et al. 2021). GridTape contains a series of apertures, each of which is covered by a plastic film. During the collection process, the sections are suspended on the films, which are transparent to electrons and hence compatible with transmission electron microscopy. This is a major difference with the original ATUM technique, which was designed to be used with scanning electron microscopy (Hayworth et al. 2014; Kasthuri et al. 2015; A. Shapson-Coe et al. 2021).

The sections were imaged by a fleet of five transmission electron microscopes. Even with the 5✕ parallelism for high speed, the microscopes operated for six months to generate over a petabyte of raw image data. The microscopes were equipped with reel-to-reel mechanisms for handling GridTape (Yin et al. 2020), enabling round-the-clock imaging that was automatically controlled by custom software.

The EM imaging data were acquired at a 4 nm pixel resolution. The axial resolution was set by the section thickness, which was nominally 40 nm. Unless otherwise noted, we employed a nominal voxel resolution of 4 ✕ 4 ✕ 40 nm^3^. Each section image was assembled by stitching together thousands of tiles, each of which contained tens of megapixels. Most stitched montages were about 60 gigapixels, though the exact dimensions varied.

A contiguous range of 19,939 sections was selected for alignment into a 3D image stack. During cutting and collection, virtually every section had developed numerous cracks and folds (Fig. 1a, 2b, 2d, S1). Points on either side of a fold had been pushed towards each other (Fig. 2a), which should be reversed for proper alignment (Fig. 2b). The image information inside the fold has been lost, so a thick black band remains after aligning a fold. The goal of alignment is not to restore the image information inside the fold, but to eliminate misalignment in the region surrounding the fold. Points on either side of a crack had been pulled away from each other (Fig. 2c), which should again be reversed for proper alignment (Fig. 2d). Some folds are wide enough to swallow up entire cell bodies, and some cracks are much wider than that. The distortions are huge relative to neurite diameters, so failing to correct misalignments around these image defects drastically lowers the accuracy of subsequent segmentation. Some defects were up to 0.6 mm long. Many tiny folds, typically just a few microns long, were observed near myelin and blood vessels. Such folds generally occurred at roughly the same location in successive sections, making alignment difficult.

**Figure 1.**
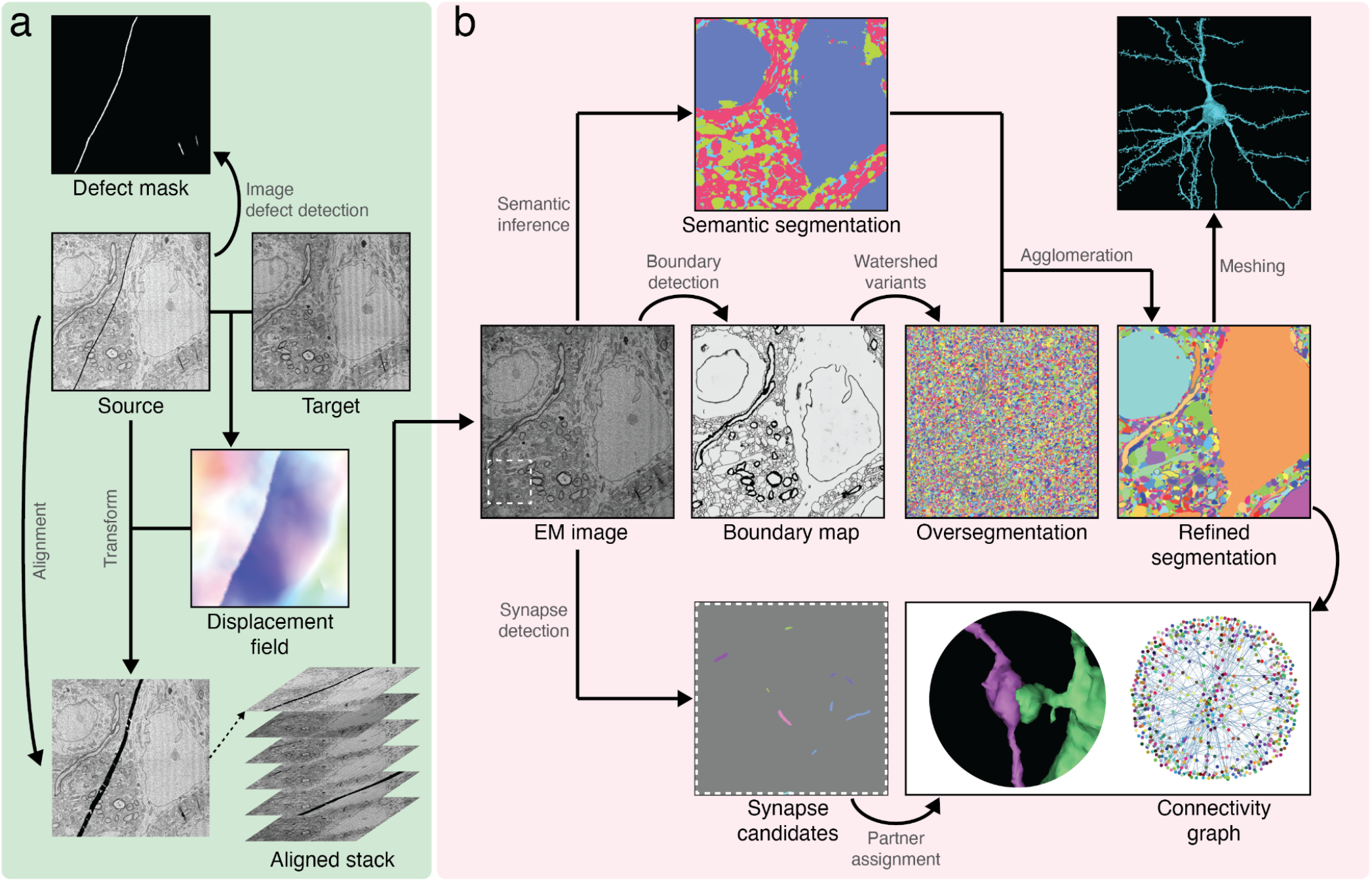
Reconstruction pipeline. (a) Image alignment and defect detection (green box). After the images were acquired, the displacement field was inferred based on consecutive images using convolutional nets to align the images. Defects such as folds and cracks were labeled by separate convolutional nets. (b) Automated segmentation and synapses (pink box). Neuronal boundaries, semantic labels (e.g. soma, axon), and synaptic clefts were detected from the EM images using convolutional nets. Neuronal boundaries were oversegmented into supervoxels by a watershed-type algorithm. Supervoxels were agglomerated with information from the neuronal boundaries and semantic labels, creating refined segments. Synaptic partners in the refined segmentation were assigned to synapses resulting in the connectivity graph of refined segments. Refined segments were meshed so they can be visualized in 3D.

**Figure 2.**
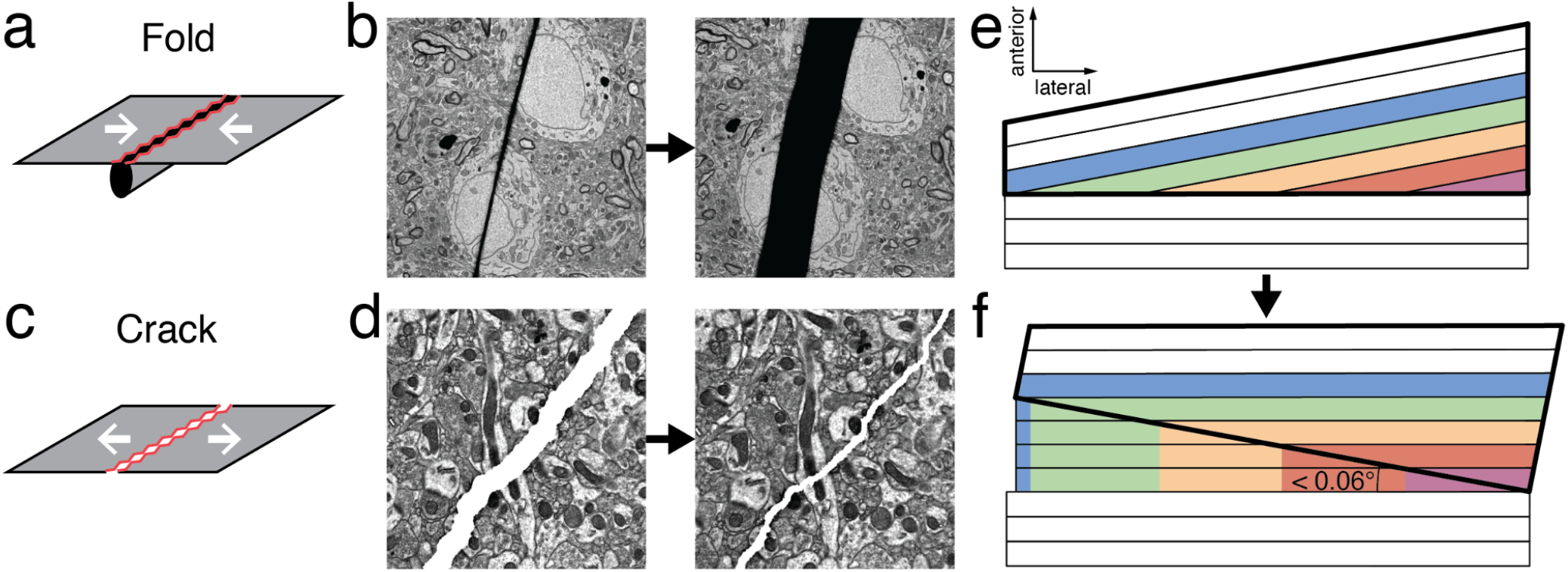
Defect handling in alignment. (a) Points on either side of a fold were pushed towards each other during section collection. (b) Alignment reverses this distortion, though a black band remains where image information was lost. (c) Points on either side of a crack were pulled away from each other during section collection. (d) Alignment reverses this distortion; the white jagged line in the right panel would be eliminated completely if alignment were perfect. (e) Schematic illustrating sectioning (bold trapezoid) following hiatus. Partial sections (colored) occurred because the new cutting plane was slightly tilted (< 0.06°) relative to the old one. The cutting sequence was bottom to top. (f) The aligned image used for reconstruction, which treats all sections as if there were no shift in the cutting plane. Each partial section was completed by compositing with larger partial sections, leading to regions of image duplication (vertical colored columns).

In a previous terascale pipeline that was built to reconstruct a pilot dataset of the MICrONS project (N. L. Turner et al. 2020; Casey M. Schneider-Mizell et al., 2020; Sven Dorkenwald et al. 2019), we used traditional computer vision methods (Saalfeld et al. 2012) supplemented by human-in-the-loop exception handling. This left misaligned bands of tissue surrounding cracks and folds. We blacked out the misaligned regions, and our automated segmentation was able to trace neurites through the missing data with fairly high accuracy (Sven Dorkenwald et al. 2019). However, the old approach was not applicable to the current dataset because it would have blacked out such a large fraction of the data due to the high frequency of image defects. Furthermore, the approach would have required prohibitive human-in-the-loop labor, because our petascale volume was 400✕ larger than the pilot dataset, and also had a much higher defect rate per unit volume.

Instead we developed a new method based on four elements: (1) a multiscale convolutional net trained by self-supervised learning for one-shot prediction of a transformation that aligns a source image to a target image (Fig. 1a) (Mitchell et al. 2019), (2) a procedure for iteratively refining the transformation by gradient-based optimization of image similarity, (3) a convolutional net that identifies image defects (Fig. 1a), and (4) a procedure for composing transformations to align a long series of images. These elements will be described in detail elsewhere; here only an overview is given.

The mean squared error between the target image and the aligned source image is used as the objective function for training the convolutional net that predicts the transformation, and also for iterative refinement of the transformation. In addition to the mean squared error, there is a penalty term that favors smooth transformations. The penalty term is gated by the defect detector, to allow nonsmooth transformations near image defects (Fig. 1a).

At inference time, the convolutional net proposes a transformation, which is then iteratively refined by gradient-based optimization. Successive transformations are composed in a way that suppresses drift. This approach removes the need for a time-consuming global elastic relaxation, facilitating the longest series of sections ever aligned and eclipsing a previous record of 10,000 sections (Morgan et al. 2016).

This dataset contained a series of partial sections that occurred after a long hiatus in sectioning (Fig. 2e, Movie S1), during which the sample was retrimmed and the knife was replaced (MICrONS Consortium et al. 2021). After the hiatus, the initial sections were partial because the new cutting plane was slightly tilted relative to the old cutting plane (Fig. 2e). During alignment, sections were processed in a sequence that was opposite from the cutting order. In particular, the partial sections were processed from largest to smallest. Each partial section was completed by duplicating pixels from larger partial sections (Fig. 2f). The smallest partial section was extended to a composite image of all the partial sections (Fig. S2). In the final aligned stack, there is a stretch of 38 partial sections that have been completed by compositing. In the “filled-in” region, neural morphologies are unnatural but traceable by human or computer. In a recent human cortex EM dataset, there were two stretches of tens of partial sections, which were treated similarly (A. Shapson-Coe et al. 2021). It seems likely that robust handling of partial sections will be a standard requirement for petascale EM reconstruction.

## Cell segmentation

Boundaries between neurons were detected by applying a convolutional net (Lee et al. 2017; Sven Dorkenwald et al. 2019) to the EM downsampled to 8 ✕ 8 ✕ 40 nm^3^ resolution rather than the original 4 ✕ 4 ✕ 40 nm^3^. Lowering the resolution economized computation and storage costs, with little effect on accuracy. Image defects such as cracks, folds, missing data, and misalignments posed challenges to boundary detection. We noticed that our networks handled sections of missing data better than several other defects when trained with appropriate image augmentations. Consequently, we identified these defects in the entire volume using separate classifiers and actively masked these locations out within the input to the boundary detection network (Methods). We also zeroed out the boundary map where there were three consecutive sections of missing data, masked input information, or remaining misalignments (Methods), because boundary detection accuracy becomes marginal at these locations. Zeroing the boundary map biases the subsequent segmentation procedure toward split errors.

A separate convolutional net was applied to semantically label voxels as (1) soma and nucleus, (2) axon, (3) dendrite, (4) glia, and (5) blood vessel (Fig. 1b). Semantic labeling has been demonstrated to improve segmentation accuracy by imposing constraints (Krasowski et al. 2018; Wolf et al. 2020; Alexander Shapson-Coe et al., 2021), to restrict segmentation to neuropil (P. H. Li et al. 2019; A. Shapson-Coe et al. 2021), or to inform automated techniques to clean an initial segmentation (A. Shapson-Coe et al. 2021; H. Li et al. 2020).

The boundary map was segmented by watershed (Zlateski and Seung 2015) and hierarchical agglomeration (Lu, Zlateski, and Seung 2021). Agglomeration was based on the mean affinity criterion (Lee et al. 2017; Funke et al. 2019), with two major modifications. Size thresholds were imposed to avoid merging big segments connected by low mean affinities. During the agglomeration process, an object was assigned to a semantic class if that semantic label predominated its voxels. Agglomeration was suppressed for a large object having weak mean affinities with objects of other semantic classes.

The volume was divided into subvolumes containing roughly 65% and 35% of the sections respectively, or subvolume-65 and subvolume-35, with the division occurring near the stretch of partial sections. The automated segmentations of the subvolumes were done separately. So far proofreading has been performed on subvolume-65 only. As mentioned earlier, the alignment is good enough to allow manual or automated tracing from one subvolume to another. Uniting the two volumes in a single automated segmentation is left for future work.

## Synapse detection and partner assignment

Synaptic clefts were automatically detected by a convolutional net (Fig. 1b) applied to the EM data at 8 ✕ 8 ✕ 40 nm^3^ resolution. The voxelwise predictions of the convolutional net were initially segmented using connected components. Each connected component was then assigned to synaptic partner segments by a separate convolutional net (Nicholas L. Turner et al. 2020), and components within a threshold distance that were assigned to the same synaptic partners were then merged to produce the final cleft detections (Methods). The final result totaled 523 million synapses (subvolume-35: 186 million, subvolume-65: 337 million).

Handling image defects in synapse detection is often less difficult than in boundary detection. Synapse detection is a highly imbalanced classification task (the “synaptic” voxels that are part of a cleft are few in comparison to the many “non-synaptic” voxels), and predicting no synapses is often a reasonable default strategy to handle defects. Consequently, no input or output masks were used for cleft detection during the prediction of the EM volume. Still, surprising and counterintuitive defects can still arise that require careful testing and dataset augmentation. As an interesting example of a rare failure mode, we mention that folds in the imaging substrate inside a blood vessel lumen (interior) after expansion during alignment were sometimes falsely detected as synapses. To handle this case, we added corrected examples of these errors to the training set and also included a filter to exclude them from the final graph, since the partner assignment step reliably predicts them to be “autapses” of the large blood vessel segments.

However, this task’s imbalance also represents a challenge in training. Human annotators often miss synapses due to errors of inattention related to this imbalance (Plaza et al. 2014), and also because it is difficult to detect clefts that are nearly parallel to the sectioning plane in serial section EM images.

Ground truth with a low recall (the percentage of true synapses captured by the labels) can make training networks with high recall difficult, especially when combined with the task’s imbalance.

Others have addressed these problems by increasing the weight of “synaptic” voxels relative to background voxels in the loss function for training (A. Shapson-Coe et al. 2021; Buhmann et al. 2021). We found it more effective to instead address the false negatives in the training labels. We improved the accuracy of the ground truth using an iterative procedure. We trained a synapse detector, and tuned inference parameters to produce few false negatives on the training set. We often found that some of the resulting “false positives” were in fact true positives that were missed by the human annotators (Nicholas L. Turner et al. 2020). These were corrected in the training set, and we iterated this procedure until no more errors of the human annotators were found.

## Ground truth annotation and augmentation

Our reconstruction pipeline included convolutional nets for boundary detection, semantic labeling, synapse detection, and synaptic partner assignment. The nets were trained using ground truth created by human annotators. We took the creation of ground truth very seriously, because it often ends up being the main determinant of real-world performance for supervised learning. Our boundary detector, in particular, required extremely high accuracy, because even a single segmentation error can lead to thousands of invalid connections in the wiring diagram. Therefore the following description is mainly focused on ground truth for boundary detection.

### Leveraging ground truth of other datasets

A larger training set generally leads to higher accuracy, but ground truth is laborious to generate. To save labor, we reused 943 μm^3^ of ground truth from pilot datasets of the MICrONS project (N. L. Turner et al. 2020; Casey M. Schneider-Mizell et al., 2020; Buchanan et al. 2021; Sven Dorkenwald et al. 2019). A boundary detector pretrained on this old ground truth turned out to generalize well to the new dataset. We further improved accuracy by continuing to train the convolutional net after adding 425 μm^3^ of ground truth from the new dataset. These datasets varied in voxel resolution and image quality. For comparison, the recent *Drosophila* hemibrain reconstruction (Scheffer et al. 2020) relied on 432 μm^3^ of manual ground truth and 31,063 μm^3^ of semi-automated ground truth to train segmentation models.

Many groups have tried semi-automated creation of ground truth, i.e., correcting boundaries of an automated segmentation (Scheffer et al. 2020). The starting automated segmentation tends to be inaccurate in the most difficult and tricky locations, which is where we most need accurate training data to improve the performance of the boundary detector. In our early experiments, we found it more straightforward to annotate such tricky locations from scratch, so we ended up sticking with purely manual annotation.

### Handling diverse cellular structures

It is relatively straightforward to increase the amount of training data. More nontrivial is the challenge of “covering the problem space with the training data” (Smith and Topin 2016). The dataset contains diverse cellular structures, and these should be thoroughly sampled by the training data (Figs. 3a, b).

**Figure 3.**
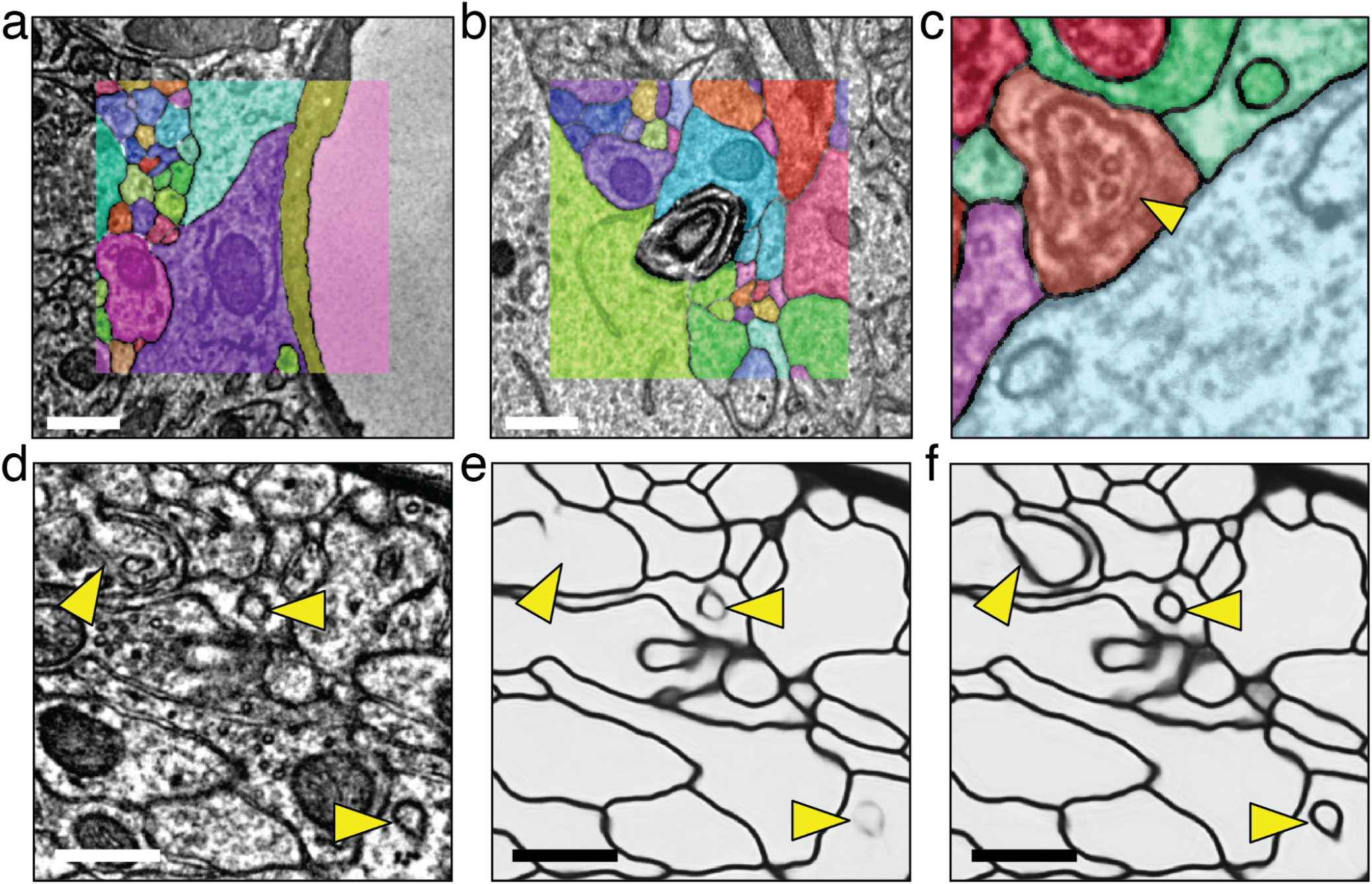
Ground truth annotation for diverse cellular structures. Example annotated volumes for ground truth of (a) a blood vessel, and (b) myelin whorl. (c) Incorrect ground truth annotation for an invaginating structure (yellow arrowhead). The invaginating structure is identifiable by clear double membranes and inclusion of synaptic vesicles, but were mistaken for intracellular organelles during labeling. (d) Examples of invaginating structures (yellow arrowheads) in EM images. (e, f) The convolutional net failed to predict correct boundaries before the ground truth correction (e), but succeeded after the ground truth correction (f). (a, b, d-f) Scale bars = 500 nm.

Sampling is complicated by the fact that the kinds of cellular structures can vary with cortical depth, and the quality of staining also varies within the imaged volume. We adopted an iterative process of ground truth creation: training a boundary detector, reconstructing a test volume, analyzing the errors to identify failure modes, and targeting new ground truth creation at the failure modes. Failure modes for segmentation included direct contact between somas, blood vessels, myelinated axons, and invaginating structures.

### Simulating image defects

Image defects are particularly pronounced in serial section EM, the method employed for the present study, compared to block face EM (Lee et al. 2019). Image defects often accompanied image distortions and loss of information. Misalignments were often caused by (and hence co-occurred near) other kinds of image defects. Therefore we found it infeasible to annotate real image defects present in the dataset to create consistent ground truth with certainty. Instead we simulated the defects by distorting clean images and their annotations. Relative to annotating real image defects, this strategy allowed us to intensify the frequency and severity of each kind of defect during simulation to improve the robustness of convolutional nets. This simulation-based approach to training data augmentation previously proven effective for EM images (Lee et al. 2017; Funke et al. 2017). The training set for the boundary detector included simulated misalignments, as was the case for a pilot dataset (Sven Dorkenwald et al. 2019; N. L. Turner et al. 2020). It was particularly important to simulate misalignments that co-occurred with missing data, which were common in the aligned stack. We also simulated partial and full missing sections, blurry (out-of-focus) sections, thicker-than-usual sections, and lost sections.

### Ground truth consistency

Besides the amount and coverage of the training data, we also learned that inconsistency in human annotations can be a significant cause of errors by a boundary detector. As a representative example, we observed systematic boundary detection failures for a variety of invaginating structures (Boyne and McLeod 1979; Boyne and Tarrant 1982; Petralia et al. 2015, 2021). The boundary detector is trained to detect the borders between cells, while ignoring intracellular membranes of organelles. In a single 2D section, an invaginating structure can resemble an intracellular organelle (Fig. 3d), and this resemblance can fool the convolutional net into “erasing” the borders of the structure, leading to a merge error in subsequent segmentation (Fig. 3e).

Upon close inspection of the ground truth, we discovered that the boundaries of invaginating structures were not consistently handled. We updated our annotation policy, and had our human annotators review all our ground truth. After retraining with the corrected ground truth, the merge errors at invaginations were substantially reduced (Fig. 3f).

## Distributed computation

Our petascale reconstruction pipeline makes extensive use of distributed and cloud computing. We processed the dataset using Google Cloud, but our pipeline can also run on other commercial clouds with little or no modification.

Because the section images are so large, alignment operations, such as generating and combining transformations, are computed for overlapping patches and blended together. The computations for the patches are distributed over many GPUs. There are minimal task dependencies. The implementation is fairly complex, and will be described elsewhere.

Boundary detection, semantic labeling, and synaptic cleft detection model inference were implemented using Chunkflow (J. Wu et al. 2021). This software package was developed previously for our terascale pipeline, and extended with only minor changes for the petascale dataset. Chunkflow divides the input image into overlapping chunks, distributes the processing of the chunks by the convolutional net across multiple workers, and recombines the results of processing into a single output image (Fig. 4a).

**Figure 4.**
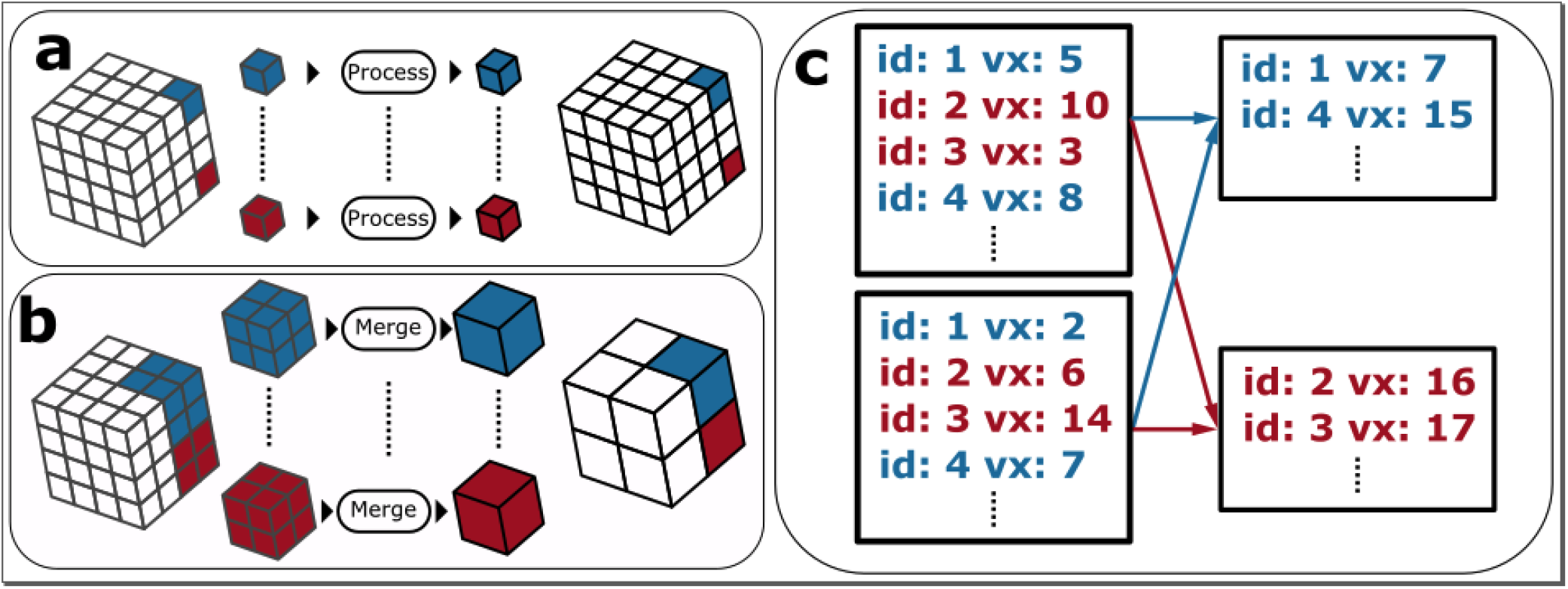
Distributed computation. The divide-and-conquer strategies employed by parts of the petascale pipeline. (a) Standard independent processing of local regions of data (“chunks”). This strategy is used by the image alignment, convolutional net inference, segmentation, and synapse processing. Each chunk may overlap with its neighbors. (b) An octree merge operation employed by the segmentation. (c) An example map-reduce operation employed by the synapse processing. Voxel counts of individual synapse predictions are summed across machines in parallel by mapping prediction ids.

Chunkflow uses Docker containers, cloud-based task queues, and Kubernetes for deployment in the cloud. This deployment is fault tolerant, robustly handling task failure through a visibility timeout mechanism. Such failures are inevitable when processing a petascale dataset, which is divided into hundreds of thousands of chunks processed by thousands of cloud GPUs running for several days (J. Wu et al. 2021). The distributed computation was simple because there were no task dependencies for different chunks.

The agglomeration step of cell segmentation (Fig. 1b) is in principle a global operation, as the mean affinity criterion summarizes information between two segments across the boundary map. We devised a way to initially parallelize agglomeration in chunks (Fig. 4a), and yet produce the same result as if agglomeration had been global (Lu, Zlateski, and Seung 2021). This was done by prohibiting agglomeration of segments near chunk boundaries. At each step of the hierarchical procedure, the chunk size was doubled in all dimensions (Fig. 4b). In a new larger chunk, some segments that had formerly been near chunk boundaries were now securely in the interior, and became eligible for agglomeration. The hierarchical procedure was recursively repeated until a single chunk encompassed the entire volume.

Hierarchical agglomeration was more complex to implement because the hierarchical task decomposition contains dependencies, and different types of tasks ran on different cloud instance types to maximize efficiency. We used Apache Airflow to manage workflow orchestration. The workflow was described by DAGs (directed acyclic graphs). The segmentation tasks were nodes in these DAGs, with the edges describing the dependencies among the tasks. The auxiliary operations that managed the cloud instance groups were also represented by nodes in the DAGs, the clusters were ramped up when the corresponding segmentation tasks were ready to run, and scaled down when the tasks were finished. Agglomeration required several weeks, sometimes using over 50,000 cloud CPUs at a time.

Synapse segmentation and synaptic partner assignment (Fig. 1b) were implemented using Synaptor. This package includes a distributed implementation of connected components that begins using chunkwise processing (Fig. 4a), and uses map-reduce processing steps to perform merging across chunks (Fig. 4c). This implementation has also been used to segment nuclei and organelles in addition to synapses (N. L. Turner et al. 2020). The package also includes an inference engine for the partner assignment convolutional net that employs both chunkwise and map-reduce steps. Container orchestration was managed using Docker containers, cloud-based task queues, and Kubernetes.

## Data access and visualization

Each step of the reconstruction pipeline described above (Fig. 1) produces several intermediate results. These intermediate results are generally images. We adopted the Precomputed format of Neuroglancer, Google’s WebGL-based viewer for volumetric data (Maitin-Shepard 2019), to view these images throughout processing, and this transparency turned out to be indispensable for debugging during pipeline development. Views of problematic locations in image stacks and their effects on the reconstruction could be exchanged effortlessly as URLs by members of the development team.

We also built the Igneous package to generate and manage Precomputed data layers. Igneous can create a multiscale image pyramid by fast downsampling of a large image, as well as generate meshes and skeletons from the cell segmentations in Precomputed format. These features collectively allowed us to diagnose failure modes in the segmentation performance in a way that would be infeasible by viewing the full-resolution segmentation image alone. Igneous generates meshes using marching cubes (Lorensen and Cline 1987) and performs mesh simplification using quadric error metrics (Garland and Heckbert 1997; Hoppe 1999). It performs skeletonization by running Kimimaro, an algorithm based on TEASAR (Sato et al. 2000; Bitter, Kaufman, and Sato 2001) that we further refined.

Igneous also handles many chunks within the large EM volumes, so it includes a task queuing mechanism inspired by ChunkFlow to ensure robustness.

Pipeline operations also communicated with each other by reading from and writing to cloud storage using CloudVolume, a serverless Python client (Silversmith 2021). Reading and writing were typically parallel and multithreaded to maximize throughput, and various options for lossless image compression were used. Of note, fpzip (Lindstrom and Isenburg 2006) was used to make the storage of the three-channel 3D floating point affinity maps generated by boundary detection practical.

Our development environment also makes public access to the results relatively straightforward. With a few careful considerations of permissions and data management, the public is able to use the same intuitive tools we’ve used for their own access and exploration. Additionally, Igneous and CloudVolume are designed to avoid dependency on vendor specific features where this is feasible to ensure they will be useful for the foreseeable future.

## Examples of neurons

The accuracy of our pipeline can be qualitatively assessed by inspecting the morphology of some example neurons. Pyramidal cells are the most common class of neuron in the cerebral cortex. Their dendrites are often reconstructed accurately enough (Fig. 5a) to allow recognition of an apical dendrite extending from the soma to the pia, and basal dendrites branching near the soma. Pyramidal axons, particularly their collaterals, are much thinner than dendrites, and consequently suffer from frequent split errors. They require significant human labor to fully proofread, as will be discussed in a companion paper.

**Figure 5.**
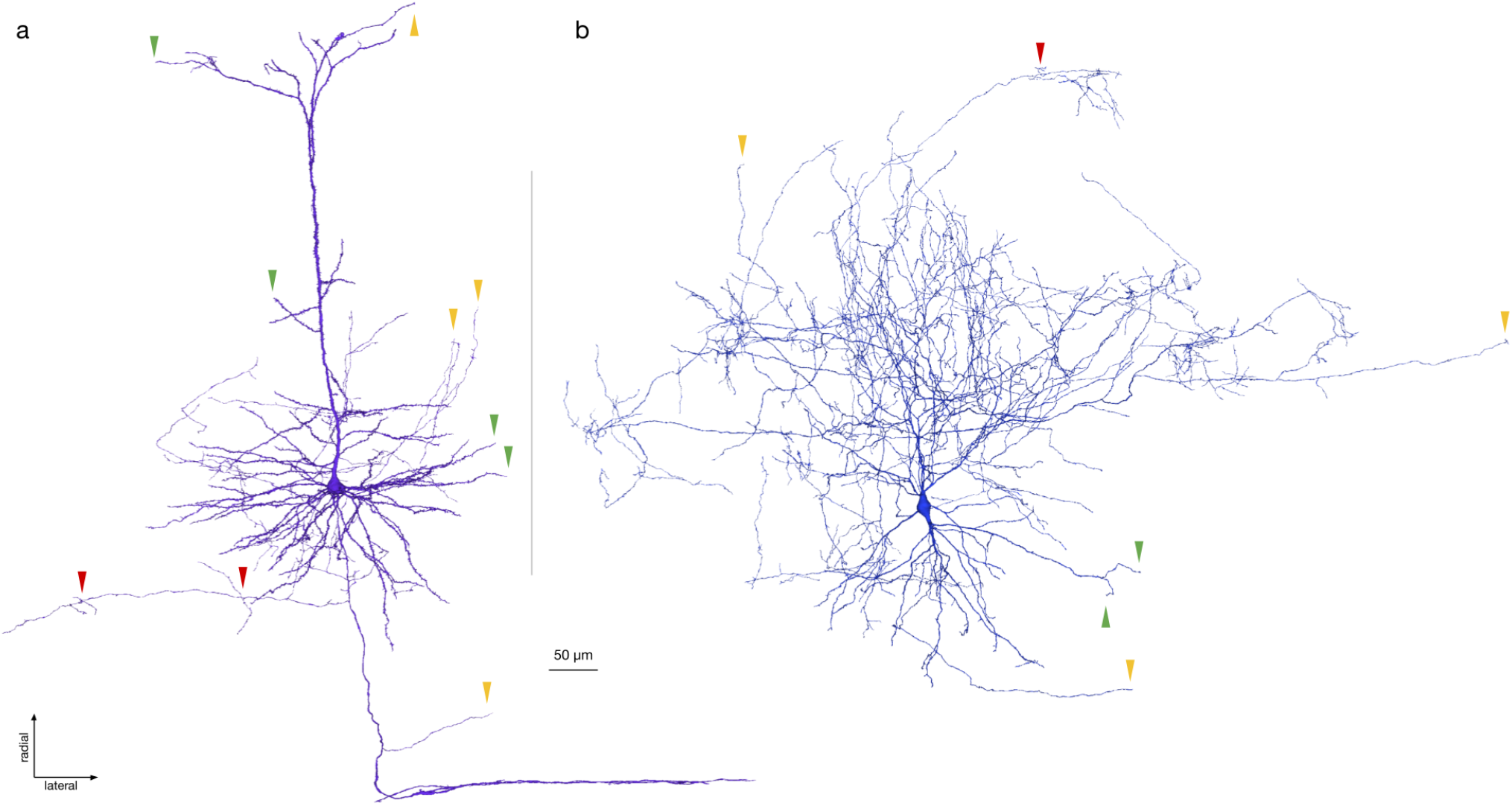
**Example automatic segmentation** of (a) L5 pyramidal cell, and (b) basket cell, projected onto the coronal plane. Arrows indicate sparsely sampled locations along a neurite that are actual terminations (green), split errors (yellow), or merge errors with another neurite (red). Largest anatomical axes after projecting onto the volume’s axes are displayed.

Neurons of certain non-pyramidal classes have thicker axons, which are often reconstructed fairly accurately. For example, the basket cell shown in Fig. 5b can be recognized by its dense axonal arbor near the soma, and sparser axonal branches hanging down below the soma into deeper cortical layers.

## Discussion

We have described a computational pipeline for reconstructing neural circuits in petascale EM datasets. The pipeline incorporates a new approach to image alignment, which was required for dealing with the high frequency of cracks and folds in the EM dataset. Most of the remaining misalignments are quite small, but misalignments are still associated with a significant fraction of segmentation errors. We optimistically predict that the frequency and size of cracks and folds will fall in future datasets, which would lead to reduction of segmentation errors associated with misalignments. According to a contrary view, serial section EM is fundamentally handicapped, and one should instead employ block face EM, a competing 3D EM technique for which alignment challenges are minimal (Lee et al. 2019a).

While more accurate ground truthing of invaginations improved segmentation accuracy, we found other configurations of cellular structures that continued to cause errors. First, sometimes a mitochondrion is so tightly close to cellular membranes that the convolutional net erroneously erases them along with the mitochondrion. Second, endoplasmic reticulum (ER) is known to occasionally form a close contact with plasma membranes (PM) (Y. Wu et al. 2017), and errors were common at such ER-PM contacts.

Besides mitochondria and ER, we observed other unknown membranous structures that are close to and often entwined with cellular membranes, causing similar problems. These errors show that it can be challenging for a convolutional net to (1) erase the membranes of intracellular organelles while (2) marking plasma membranes, a tension that is intrinsic to the problem of neuronal boundary detection. Possible solutions may include (1) learning an object-centered representation (Lee et al. 2021; Sheridan et al. 2021) to better understand the scene, and (2) conservatively detecting all biological membranes to oversegment intracellular structures and later postprocessing them, possibly with the help of biological priors (Krasowski et al. 2018).

Other errors are associated with low image contrast and limited axial resolution, especially at locations where sections are missing or thicker. Achieving high-contrast staining throughout a millimeter-scale sample is still an elusive goal, despite progress in this direction (Mikula and Denk 2015; Hua, Laserstein, and Helmstaedter 2015). Overall, computational advances in neural circuit reconstruction have led to a keen need for improved EM sample preparation and image acquisition.

Methods for automated error detection and correction would be helpful for improving segmentation accuracy. Research on such methods has demonstrated promising results (Zung et al. 2017; Rolnick et al. 2017), but has not yet been deployed at scale in our pipeline.

EM-based reconstruction of an entire mouse brain has been proposed as a transformative project for neuroscience (Abbott et al. 2020). This would be an exascale dataset, roughly 1000✕ larger than the present one. We expect that most of our pipeline operations should scale to an entire mouse brain, because the tasks for the image chunks have little or no dependencies. The primary exception is hierarchical agglomeration, which proceeds by recursive octree merging (Fig. 4b). The last step of the recursion at the top of the octree requires a shared memory machine with RAM and CPUs that scale roughly as the square of the linear size of the EM volume. In other words, the shared memory machine for the whole mouse brain would need to be 100✕ larger than the one used for the current cubic millimeter volume. This factor could be reduced with a less naive scaleup of hierarchical agglomeration, which is an interesting topic for future research.

## Supplementary figures

**Supplementary Figure 1.**
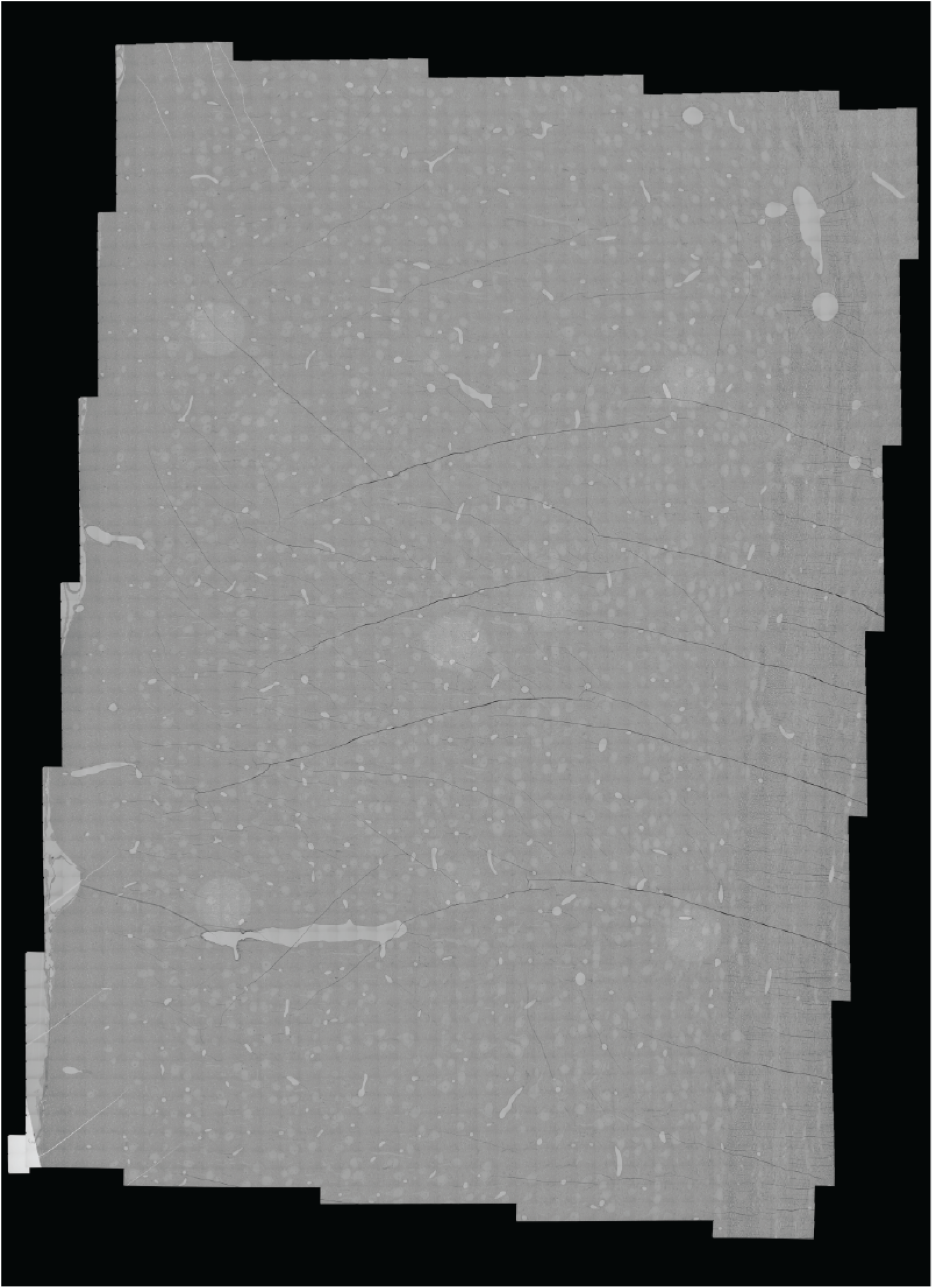
Example section (z=21014) with folds and cracks. Numerous folds and cracks exist in nearly every imaged section.

**Supplementary Figure 2.**
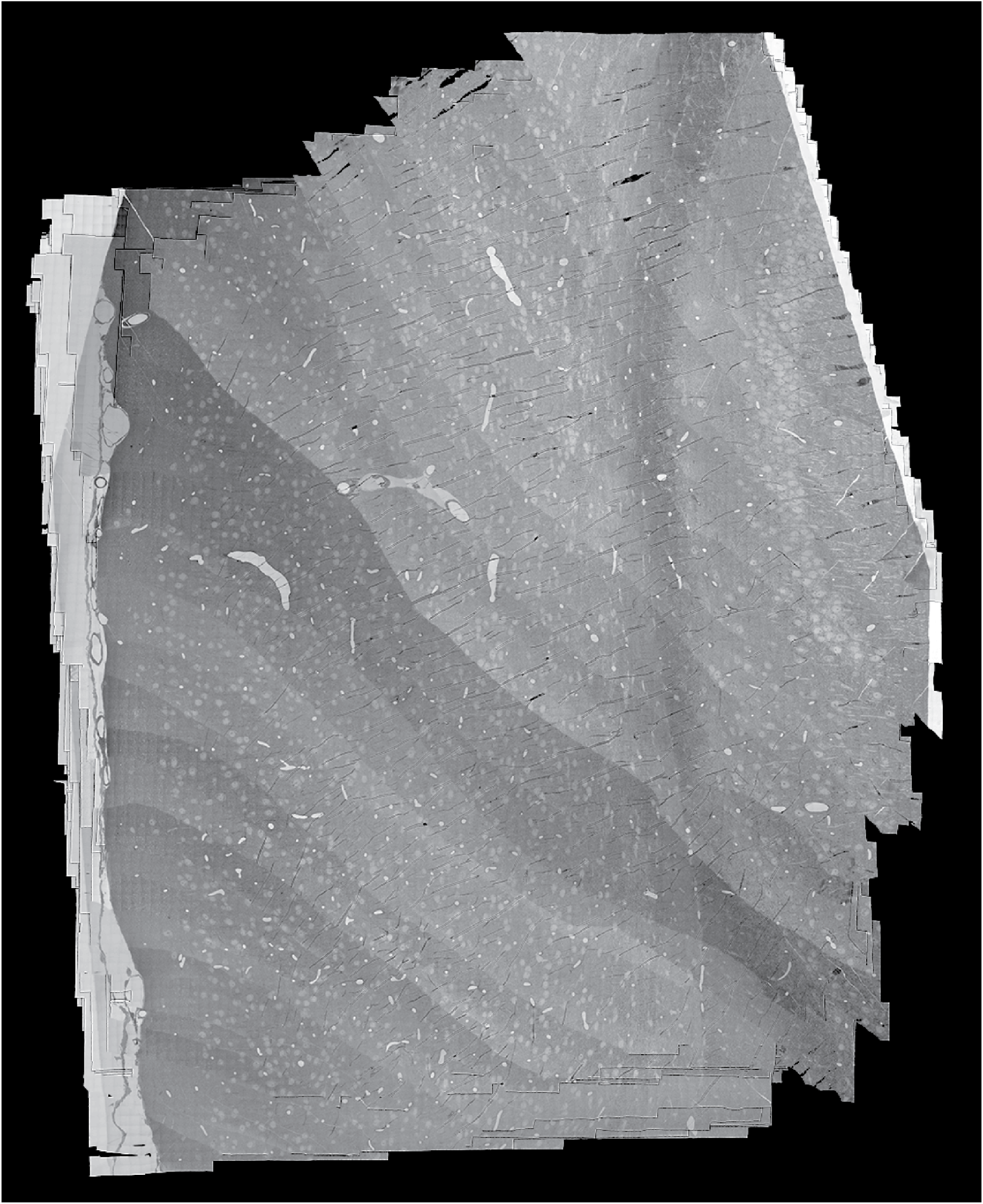
Composited image of partial sections.

## Methods

### Image acquisition

EM image acquisition and stitching followed the methods described in (Yin et al. 2020). Each section image was assembled by stitching together an average of 5,552 tiles, each of either 3,840 ✕ 3,840, 5,408 ✕ 5,408, or 5,504 ✕ 5,504 pixels. Most stitched montages were 296,000 ✕ 200,000 pixels, though these dimensions varied. Stitching software and algorithms from (Khairy, Denisov, and Saalfeld 2018) were scaled up to our large section images (Yin et al. 2020).

### Image alignment

#### Rough alignment

Each image tile represented a small region of a serial section. The tiles were first corrected for lens distortion effects. A non-linear transformation was computed for each section using a set of 10✕10 highly overlapping images collected at regular intervals during imaging. The lens distortion correction transformations represent the dynamic distortion effects from the TEM lens system and hence require an acquisition of highly overlapping calibration montages at regular intervals. Overlapping tile pairs were identified within each section, and point correspondences were extracted for every pair using SIFT feature descriptors. Per tile transformation parameters were estimated by a regularized solver algorithm that minimized the sum of squared distances between the point correspondences between these tile images. Deforming the tiles within a section based on these transformations resulted in a seamless stitching of the section.

A downsampled version of these stitched sections was produced for estimating a per-section transformation that roughly aligned these sections in 3D. A process similar to 2D stitching was followed here, where the point correspondences were computed between pairs of sections that were within a desired distance in z direction. The per-section transformation was then applied to all the tile images within the section to obtain a rough aligned volume. MIPmaps were utilized throughout the stitching process for faster processing without compromise in stitching quality.

#### Large crack correction

34 sections contained severe cracks (displacements greater than ~30 μm), which we decided to manually correct. Using Neuroglancer (Maitin-Shepard 2019), point-pair correspondences were manually made between each of the following sections and a neighboring section:

15016, 15022, 15070, 16001, 17448, 18028, 19713, 21560, 23992, 24424, 24425, 24860,
25333, 25509, 25756, 25849, 26020, 26043, 26124, 26158, 26230, 26236, 26237, 26440,
26579, 26698, 27088, 27089, 27092, 27171, 27210, 27211, 27246, 27765

We computed the Delaunay triangulation for the correspondence vertices in each affected section, and a piecewise affine model per triangle was used to transform each section at 8 ✕ 8 ✕ 40 nm^3^ resolution before proceeding to later alignment stages.

#### Fold & crack detection

We trained separate convolutional networks to detect cracks and folds by producing voxel-wise predictions on the rough aligned data at 64 ✕ 64 ✕ 40 nm^3^ resolution. The network architectures were identical 2D U-Nets. The architecture consisted of three 3 ✕ 3 non-strided convolutions followed by 4 down convolution layers with max pooling and three 3 ✕ 3 non-strided convolutions, 4 up convolution layers with nearest-neighbor interpolation and three 3 ✕ 3 non strided convolutions, and one output layer with one 3 ✕ 3 non-strided convolution. The feature map numbers were [16, 32, 64, 128, 256] from the first layer to the deepest layer respectively. The network operated on a 256 ✕ 256 px field of view (Ronneberger, Fischer, and Brox 2015). The fold training set contained 3,830 samples of image and label pairs, while the crack training set contained 3,894 samples. The samples had size of 256 ✕ 256 px. Labels were manually annotated with VAST (Berger, Seung, and Lichtman 2018). Training augmentations included (1) flip & rotate and (2) brightness & contrast augmentations described in the Dataset Augmentation section below. Inference was distributed in chunks of 2048 ✕ 2048 across each section with 50% overlap between adjacent chunks in both X and Y.

#### Resin detection

We trained a convolutional network to detect resin. Voxel-wise predictions were made on the rough aligned data at 2048 ✕ 2048 ✕ 40 nm^3^ resolution. The network architecture and dataset augmentation was identical to the fold and crack detection. The training set included 50 samples of image and manually labeled pairs.

#### Alignment at 1024 nm & 64 nm

Using the method outlined in (Buniatyan et al. 2020), we trained a convolutional network to produce an embedding for each section to improve accuracy during patch matching. We embedded the 1024 ✕ 1024 ✕ 40 nm^3^ sections after masking voxels identified as resin to zero. We computed displacement fields between neighboring embedded sections (up to five previous sections) by cross-correlating a co-located 256 x 256 px source patch with a 384 x 384 px target patch at each vertex in a 96 px Cartesian grid. We combined these pairwise displacement fields to generate a displacement field for each section.

Using methods similar to those outlined in (Mitchell et al. 2019), we trained a convolutional network through self-supervision on the entire rough aligned dataset to predict a displacement field between a pair of 1024 ✕ 1024 ✕ 40 nm^3^ sections after embedding as described above, and also took as input the most recently predicted displacement field and crack and fold masks. We augmented the data with transposing image pairs, as well as translating one image relative to the other up to 4 px at this resolution. We used this network to compute displacement fields between all neighboring sections (up to five previous sections). At inference time, the predicted displacement fields were further refined by using gradient descent to optimize the similarity between the embeddings that were the input to the network. We then combined these pairwise displacement fields by ascending section index to generate a new displacement field for each section.

Similar to the above procedure, we also trained a convolutional network to produce displacement fields on 64 ✕ 64 ✕ 40 nm^3^ sections. These images were not embedded in advance. Specifically, the network was trained on a subset of the rough aligned dataset to predict a displacement field between a pair of 64 ✕ 64 ✕ 40 nm^3^ sections, transforming the highest-level encodings with the most recently predicted displacement field, and with cracks and folds masks used to adjust the similarity and smoothness penalties in the loss function. The subset consisted of a small initial group of 10 manually chosen samples that contained fold and crack patterns deemed to be particularly difficult, as well as an extended set of 1200 randomly chosen samples within the rough aligned dataset. Each sample covered 5 consecutive sections, allowing for up to 20 permutations per sample. We used this network to compute displacement fields between all neighboring sections. We then created composite sections of neighbors if more immediate neighbors were deemed to have displacements larger than 8 px relative to previous neighbors (evaluated with cross-correlation), or if they contained zero-value pixels. At inference time, the predicted displacement fields were further refined by using gradient descent to optimize the similarity between the lowest-level embeddings produced by the network. We then combined these pairwise displacement fields by ascending section index to generate a new displacement field for each section. We considered the output of this step to be the final displacement field. We bilinearly upsampled the field to 8 ✕ 8 ✕ 40 nm^3^, and transformed the rough aligned dataset into the final aligned dataset.

#### Alignment in two parts for handling partial sections

The data were aligned in two separate stages to handle the partial sections caused by the sectioning restart. We first aligned subvolume 65 without partial sections (sections 14828 to 27882) at both resolutions mentioned above. We then aligned across the partial sections in descending order of section index, from 14828 to 14783. The partial sections shrink with decreasing index, implying that we could best align the highest index section to the adjacent data. The last seven partial sections separating the two subvolumes either only contained small fractions of tissue exclusively outside the region of interest, or had been intentionally left blank in the unaligned stack to indicate potential tissue loss. In the aligned stack, these sections have been dropped and the full sections of subvolume 35 accordingly moved up. We created an intermediary composite dataset of the aligned partial sections that we used for reconstruction: for each aligned partial section, we created an image that sets each pixel to the value of the first non-zero pixel in the ordered set of sections from the current section to the highest-indexed partial section. We then aligned subvolume 35 in descending order of section index, using the first section in this composite dataset as the initial section. When aligning subvolume 35, we applied a bandpass filter for knife chatter artifacts oriented at 87° to the sections before creating composite images of neighbors.

### Segmentation masks

To avoid significant segmentation artifacts, we created an input mask for regions of the data remaining misalignments or without useful image content. This mask sets the input to boundary detection to zero in these regions. We also generated an output mask that set the output of boundary detection to zero in other regions. We created these masks by evaluating locations in the aligned dataset which were still misaligned, or did not contain tissue.

We produced misalignment maps between each section and its neighbors, up to five sections apart, for both the 256 ✕ 256 ✕ 40 nm^3^ aligned dataset, as well as 1024 ✕ 1024 ✕ 40 nm^3^ aligned dataset. To compute a misalignment map for a pair of images, we smoothed a thresholded correlation of 8 ✕ 8 px patches that have been transformed and high-passed filtered in the frequency domain. We used the misalignment maps to classify misalignments as either “step” misalignments (pairs of two consecutive sections that are not aligned to each other, but predecessors are aligned to the first section in the pair and successor sections are aligned to the second section of the pair) or “slip” misalignments (single sections that are neither aligned to its predecessor nor its successor, but both predecessor and successor are aligned to each other).

An input mask for segmentation removed all slip misalignments, folds, cracks, and resin masked voxels from the input to the boundary detection network. The output mask removed all step misalignments, as well as any location masked out of the input mask for three consecutive sections.

### Boundary Detection

Using the method outlined in (Lee et al. 2017), we trained a convolutional network to estimate inter-voxel affinities that represent the potential for neuronal boundaries between adjacent image voxels. The network was pretrained using manually annotated ground truth from previously collected cortical datasets (N. L. Turner et al. 2020; Sven Dorkenwald et al. 2019; Buchanan et al. 2021; Casey M. Schneider-Mizell et al., 2020), then finetuned on a mix of ground truth collected from this dataset, as well as the previous cortical datasets. We made modifications in network architecture, auxiliary tasks, and data augmentation during training as follows.

#### Network architecture and auxiliary tasks

We used a modified version of “Residual Symmetric U-Net’” of (Lee et al. 2017). First, we added an extra convolution layer to every “residual module” (Lee et al. 2017). Second, we used 3 ✕ 3 ✕ 3 convolution everywhere except for the input and output convolution layers which use 5 ✕ 5 ✕ 1 filters. Third, we replaced exponential linear units (ELUs, (Clevert, Unterthiner, and Hochreiter 2015)) by rectified linear units (ReLUs). Fourth, we used Instance Normalization (Ulyanov, Vedaldi, and Lempitsky 2016) instead of Batch Normalization (Ioffe and Szegedy 2015).

Besides nearest neighbor and long-range inter-voxel affinities (Lee et al. 2017), we additionally detected (1) myelin and (2) blood vessel walls (endothelial cell lining) as auxiliary training targets. Explicit detection of blood vessel walls helped reduce failures in affinity prediction at blood vessel boundaries. However, we found that the two additional auxiliary tasks slightly impede the optimization of the main affinity loss. Therefore, we used both of these tasks in pretraining, but discarded the detection head for blood vessel wall during finetuning. We kept the myelin detection head during finetuning because we planned to use the myelin detection result for postprocessing affinity maps.

Overall, the net takes as input a 256 ✕ 256 ✕ 20 voxel patch at the voxel resolution of 8 ✕ 8 ✕ 40 nm^3^, and produces (1) nearest neighbor affinity maps (3 channels for x, y, and z-direction, respectively), long-range affinity maps (9 channels), (3) a probability map for myelin (1 channel), and (4) a probability map for blood vessel walls (1 channel). Each output volume has the same size as as the input. For long-range affinities, we used (x,y,z) offsets of (−5,0,0), (−10,0,0), (−15,0,0), (0,−5,0), (0,−10,0), (0,−15,0), (0,0,−2), (0,0,−3), and (0,0,−4). At inference time, we discarded the long-range affinities and only generated the nearest neighbor affinities and myelin predictions, with the input inference patches of 256 ✕ 256 ✕ 20 voxels at the voxel resolution of 8 ✕ 8 ✕ 40 nm^3^.

#### Loss function

For affinity prediction, we used the binary cross-entropy loss with “inverse margin” of 0.1 (Huang and Jain 2013). This loss function is the same as the standard binary cross-entropy loss except that targets are 0.9 and 0.1 instead of 1 and 0 respectively. Such “soft targets” are thought to have some regularization effect by penalizing overly confident predictions (Huang and Jain 2013). Although the use of soft targets resulted in slower convergence (a typical consequence of regularization), we found no conclusive evidence for better generalization.

We also used the patch-wise class rebalancing as in (Lee et al. 2017) to compensate for the lower frequency of boundary voxels. We computed in each training patch the number of positive and negative examples (boundary and non-boundary voxels, respectively), and weighted each example (or voxel) inversely proportional to the number of the class (positive/boundary or negative/non-boundary) it belongs to. Unlike the soft targets, we found that class rebalancing is critical for reducing false negative detection (missing boundaries) and thus improving segmentation quality after postprocessing.

For myelin detection, we used the same binary cross-entropy loss with inverse margin of 0.1, but no class rebalancing was used. The same loss was used for blood vessel detection during pretraining.

#### Training details

Dataset augmentation included (1) flip & rotation, (2) brightness & contrast perturbation, (3) warping, (4) misalignment, (5) missing section, (6) out-of-focus section, (7) misalignment + missing section, (8) lost section, (9) lost + missing section, and (10) slip interpolation. See the dataset augmentation section below. Misalignment augmentation used three different severities of misalignment: (1) *small misalignment* with maximum displacement *d*_*max*_ of 10 voxels in x/y-dimension, (2) *medium misalignment* with *d*_*max*_ of 30 voxels in x/y-dimension, and (3) *large misalignment* with *d*_*max*_ of 50 voxels in x/y-dimension. For simulated misalignments, we chose “step” type misalignments with *p* = 0. 7and “slip” type with *p* = 0. 3 (Lee et al. 2017).

We trained our nets on four (pretraining) or eight (finetuning) NVIDIA Titan X Pascal GPUs using synchronous gradient update. We used the AMSGrad variant (Reddi, Kale, and Kumar 2019) of the Adam optimizer (Kingma and Ba 2014), with α = 0. 001, β_1_ = 0. 9, β_2_ = 0. 999, ε = 10^−8^. We used a single training patch (minibatch of size 1) for each model replica on GPUs at each gradient step. We pretrained our nets for 550K iterations, and finetuned for 1M iterations.

### Semantic labeling

Segments within the manually segmented ground truth were viewed and classified into one of 9 semantic categories over 18 training volumes. The 9 semantic categories were (1) soma, (2) axon, (3) dendrite, (4) glia, (5) blood vessel, (6) endothelial cell, (7) myelin sheath, (8) nucleus, and (9) unknown. For the purposes described below, we computationally fused the labels for soma and nucleus, and removed the unknown, endothelial cell, and myelin sheath labels from training. This preserved 5 target classes for prediction: (1) soma+nucleus, (2) axon, (3) dendrite, (4) glia, and (5) blood vessel.

These original training volumes represented 333.7 μm^3^ of tissue. These were split into a 253.5 μm^3^ training set and a 80.2 μm^3^ validation set. After preliminary experiments, we produced two additional volumes of sparse ground truth to improve blood vessel prediction (5.6 μm^3^ and 8.3 μm^3^) from tests of the segmentation workflow. These sparse volumes only labeled automated predictions of blood vessels, and we removed the other automated segments from training.

A convolutional net was trained to perform a semantic segmentation of the EM image. The trained model takes 3D patches of the EM image as input and produces five 3D volumes as output. Each volume represents a target class specified above. Within a given output volume, the desired output for each voxel is a binary classification predicting the presence of that volume’s semantic class within the EM image.

The network architecture is a version of the Residual Symmetric U-Net (Lee et al. 2017) with six modifications. First, the number of upsampling and downsampling layers were increased to 5 along with the number of feature maps (32, 64, 128, 256, 512 at each downsampling level). Second, 3 ✕ 3 ✕ 3 convolution filters were used everywhere except for the input and output convolution layers, where 5 ✕ 5 ✕ 1 filters were used. Third, we used upsampling by nearest neighbor interpolation. Fourth, we used Instance normalization (Ulyanov, Vedaldi, and Lempitsky 2016) instead of Batch Normalization (Ioffe and Szegedy 2015). Fifth, we added an extra convolutional layer with filter size of 1 ✕ 1 ✕ 1 at the output layer. Finally, the input patch size to the neural network was fixed to 224 ✕ 224 ✕ 18. The output patch size for each output volume was the same size as the input patch.

We used a binary cross entropy loss on each output volume to train the network. We augmented dataset samples using (1) flip & rotate, (2) brightness & contrast, (3) misalignment, (4) missing section, and (5) misalignment + missing section techniques described in the Dataset augmentation section.

Misalignment augmentation used three different severities of misalignment: (1) *small misalignment* with maximum displacement *d*_*max*_ of 5 voxels in x/y-dimension, (2) *medium misalignment* with *d*_*max*_ of 15 voxels in x/y-dimension, and (3) *large misalignment* with *d*_*max*_ of 25 voxels in x/y-dimension. For simulated misalignments, we chose “step” type misalignments with *p* = 0. 7 and “slip” type with *p* = 0. 3.

We performed all experiments with PyTorch. We trained the network on NVIDIA GeForce GTX 1080Ti having approximately 11GB of memory. We used Adam optimizer (Kingma and Ba 2014) with a fixed initial learning rate of 1 ✕ 10^−5^ and trained it for 300 thousand iterations (approx 4 days). After this initial phase, we continued training the best performing state of the model as measured on the validation set using the Intersection over Union (IoU) metric. We continued training this state for a further 300 thousand iterations with a decreasing learning rate. The learning rate was decreased by 10 percent at any point during training if the validation IoU did not increase for the previous 5 thousand iterations.

### Watershed

We first processed the affinity map with a distributed watershed and clustering algorithm to produce an over-segmented image, where the watershed domains are agglomerated using single-linkage clustering with size thresholds (Lu, Zlateski, and Seung 2021; Zlateski and Seung 2015). We call the segments in this image “supervoxels”. Processing occurred in 10 million 256 ✕ 256 ✕ 512 vx chunks at 8 ✕ 8 ✕ 40 nm^3^ resolution, and combined using a 11-layer octree. These parameters were chosen to limit the surface area between the chunks being merged near the top layer of the octree. Due to the anisotropic voxel size, the contact surface in the x-y plane is much larger than the x-z and y-z planes. Starting 256 ✕ 256 ✕ 512 vx chunks makes sure we only stitches chunks along the x-z and y-z planes.

### Agglomeration

Using the method described in (Lu, Zlateski, and Seung 2021), we processed the supervoxel and affinity images to produce an region adjacency graph of the supervoxels, using a constrained mean-affinity agglomeration to produce groupings of supervoxels. Processing occurred with the same chunking parameters as watershed. The constraints in mean-affinity agglomeration are activated when mean affinities between segments are lower than 0.5. They included using the semantic label predictions to prevent neuron-glia mergers as well as axon-dendrite and axon-soma mergers. Labels were used if an agglomerated object contained at least 170 thousand voxels and at least 60% of its voxels share the same label. The constraints also included a size threshold that limited “dumbbell” type mergers joining two objects with somata. If two objects both contain more than 1000 supervoxels and one of them has more than 10 thousand supervoxels, they will not be merged.

### Synapse detection and synaptic partner assignment

We selected an initial set of 15 training volumes from initial aligned subregions of the full dataset at 8 ✕ 8 ✕ 40 nm^3^ resolution, and had human annotators label synaptic clefts within each volume. These volumes were selected to sample failure modes of other datasets - myelin, blood vessels, nuclear membranes, and organelles within somata. We added a region of “padding” to each dataset in order to give human annotators sufficient context to resolve ambiguities without checking the source images. The labeled portions of each volume constituted 227.6 μm^3^ of image data with 135 synaptic clefts.

We trained a convolutional network to predict whether a given voxel within each dataset was “synaptic” or “non-synaptic” in the sense that it was labeled to be part of a synaptic cleft. We used a Residual Symmetric U-Net Architecture (Lee et al. 2017) with 32, 40, and 80 feature maps at the different levels of the U-Net hierarchy, ReLU nonlinearity, and Instance Normalization (Ulyanov, Vedaldi, and Lempitsky 2016) instead of Batch Normalization (Ioffe and Szegedy 2015).

An initial convolutional network model was trained to evaluate the ground truth labels, using an Adam optimizer (Kingma and Ba 2014), and binary cross entropy loss function with a single static learning rate (1 ✕ 10^−4^) for 375 thousand iterations. We both edited and augmented the training set with this model’s output. Specifically, we reviewed the labeled region of each volume in comparison with the predicted clefts, and made corrections where necessary. In addition, we also augmented the training set by predicting the padding region of context, and referencing the image data surrounding the image volumes where cases were ambiguous. We also generated output for other regions of the dataset to identify any notable failure modes, and created 10 extra smaller volumes to increase training set representation of synapses onto soma among other errors. We finally added one training volume to decrease network predictions within the interior of blood vessels. This process quadrupled the size of the training set to 914.0 μm^3^ of image data, and 561 total clefts.

After finalizing the ground truth, we trained a separate network to produce output on the larger volume. For this training, we used a manual learning rate procedure, decreasing the learning rate by a factor of 10 at apparent convergence of the smoothed validation set error curve. We started training with a learning rate at 1 ✕ 10^−4^, and decreased it 5 times. As before, training used a binary cross entropy loss function. Unlike the data evaluation experiments, the training data was augmented using (1) flip & rotate, (2) missing section, (3) warping, (4) brightness and contrast, and (5) misalignment techniques described in the Dataset augmentation section below. Misalignment augmentation used one severity of misalignment with *d*_*max*_ of 10 voxels in x/y-dimension, and we chose “step” type misalignments with *p* = 0. 7and “slip” type with *p* = 0. 3. We stopped training after 1.23 million training updates.

We then trained a separate network to perform synaptic partner assignment by producing the voxels of the synaptic partners given the synaptic cleft as an attentional signal (Nicholas L. Turner et al. 2020). This used 1660 μm^3^ of total training data with 867 clefts, including data from other EM volumes (N. L. Turner et al. 2020; Sven Dorkenwald et al. 2019; Casey M. Schneider-Mizell et al., 2020; Buchanan et al. 2021). We also used the Residual Symmetric U-Net architecture for this task, using a model with 16, 20, and 40 feature maps, Instance Normalization (Ulyanov, Vedaldi, and Lempitsky 2016), an Adam optimizer (Kingma and Ba 2014), and a binary cross entropy loss with a static learning rate of 1 ✕ 10^−3^ for 2.71 million training updates. Training augmented the data using (1) flip & rotate, (2) missing section, (3) warping, (4) brightness & contrast, and (5) misalignment techniques described in the Dataset augmentation section below. This model produced perfect assignment results within our test set of 81 clefts (with 96% accuracy within the validation set of 110 clefts).

We evaluated our final cleft detection model by analyzing the overlap between the connected components of predictions and the labels. Matching how we planned to process the larger volume, we merged connected components together which had centroid coordinates within 700 nm from one another and were assigned to the same synaptic partners in each training volume. This yielded cleft detection performance estimates of 98.2 precision and 98.2 recall for our small volume test set with 56 clefts.

Following the methods described in (J. Wu et al. 2021), we predicted synaptic clefts on the entire aligned dataset. The input and output patch size of the convolutional network was 192 ✕ 192 ✕ 18. The image chunk overlap and patch overlap were both 26 ✕ 26 ✕ 4. The input image chunk size was 2516 ✕ 2516 ✕ 284, and the the output synaptic cleft map size for a single task in the cloud was 2464 ✕ 2464 ✕ 276.

We made discrete predictions of synaptic clefts by performing connected components throughout the large volume. We performed connected components in 2048 ✕ 2048 ✕ 256 voxel chunks, and then ran additional processing to merge the predictions from these chunks together. These connected components were filtered by an initial size threshold at 40 voxels (at 8 ✕ 8 ✕ 40 nm^3^ resolution). During this step, we also computed bounding boxes, sizes, and centroid coordinates for each predicted cleft. Our infrastructure used a kubernetes cluster with around 650 compute nodes for around 24 hours, and approximately 100 nodes for another day to consolidate the chunk-wise processing.

We then assigned each connected component predicted by the synapse detection within each 2048 ✕ 2048 ✕ 256 chunk of the volume to synaptic partners. This step took around 2.5 days using 800 compute nodes. Some cleft predictions crossed chunk boundaries, yet the assignment network yielded a single pair of predicted partners for interior clefts. For a cleft that crossed at least one boundary, we assigned it the synaptic partners inferred by the network within the chunk that contained the most voxels of that cleft. Similar to our small-volume evaluation, we also merged connected components together which had centroid coordinates within 700 nm from one another and were assigned the same synaptic partners. These final consolidation tasks took about 6 hours of processing over approximately 100 nodes.

### Dataset augmentation

Each convolutional network was trained using a set of dataset augmentations to improve the network’s robustness to imaging artifacts and other variability in the image data. Below we detail the set of data augmentations used across each network described above.

- **Flip & rotation**

- Each input patch to the convolutional net is randomly flipped in x and y dimensions, and then rotated by multiples of 90 degrees. The random combinations of rotation by 90° and horizontal/vertical flips are only applied in the xy-plane due to the voxel anisotropy of the EM images. This procedure yields eight 2D configurations. Additional flip in z-dimension doubles the number of configurations, resulting in a total of 16 configurations. This augmentation is commonly used in image processing applications.
- **Brightness & contrast**

- We perturbed grayscale intensity values of the input images by randomly adjusting brightness and contrast. We adapted the code from ElektroNN (Sven Dorkenwald et al. 2017), an open source high-level library for training convolutional nets (http://elektronn.org/). This is another widely used data augmentation in image processing applications.
- **Warping**

- We randomly warped the input images by combining the following linear transformations: continuous rotation, (2) shear, (3) twist, (4) scale, and (5) perspective stretch. Again we adapted the code from ElektroNN (Sven Dorkenwald et al. 2017).
- **Misalignment** (Lee et al. 2017)

- We simulated misalignments with varying severity specified by a maximum displacement *d*_*max*_. The displacement in each dimension is randomly sampled from an integer-valued discrete uniform distribution in [0,*d*_*max*_]. For each simulated misalignment, we randomly chose the type of misalignment (“slip” vs. “step”, see the Segmentation Masks section) with a predefined probability.
- **Missing section** (Lee et al. 2017)

- We simulated (1) two consecutive missing sections, either partial or full (randomly chosen), and (2) up to seven independent single missing sections, either partial or full (randomly chosen). As a result, each training patch can have at least two consecutive missing sections and no more than nine missing sections in total.
- **Out-of-focus section** (Lee et al. 2017)

- We simulated up to seven independent single out-of-focus sections by blurring the image with a gaussian kernel. We chose randomly whether to apply the blurring to the full input section, or only part of it.

Listed below are novel types of data augmentation we have newly added.

- **Misalignment + missing section**

- (Lee et al. 2017) simulated both misalignments and missing sections independently of each other. However, we found a critical flaw that can happen occasionally when the two types are independently simulated; If the two types are coincidentally simulated at the same z-section, then the simulated missing section unintentionally obfuscates the exact location (above or below the section?) of the simulated misalignment. Such ambiguity confuses the net during training, forcing it to make its best guess by predicting “double’” boundaries, each corresponding to the section above and below.
- To resolve such problematic cases, we introduce a novel *misalignment + missing section* data augmentation. Specifically, we co-simulate both types at the same location and force the net to smoothly “interpolate” through the co-occurrence of the misalignment and missing section(s). Additionally, we modified our data augmentation logic such that simulation of (1) misalignment, (2) missing section, and (3) their co-occurrence are all mutually exclusive.
- **Lost section**

- Some adjacent sections are poorly correlated with each other, meaning the content of images changes rather abruptly and non-smoothly between consecutive sections. Such non-smooth consecutive sections can result from either (1) sections entirely lost during the imaging process or (2) imperfect cutting that yields a thicker-than-normal section.
- To make our nets more robust to such non-smooth consecutive sections, we introduced a new type of data augmentation for simulating them during training. *Single lost section* removes a single image section in the input and a corresponding target section in the ground truth segmentation together from the training example. *Double lost section* removes two consecutive image sections in the input and corresponding target sections in the ground truth segmentation together from the training example. We randomly simulated both types independently during training. As a result, each training patch can have at least one lost section and no more than three lost sections in total.
- **Lost + missing section**

- We additionally introduced the *lost + missing section* augmentation. Here we replace three consecutive image sections in the input with a single missing section, and leave only the middle of the three corresponding target sections in the ground truth segmentation, so that a net can learn to interpolate the missing section as smoothly as possible in the challenging co-occurrence of missing and lost sections. To avoid any problematic situation, we modified our data augmentation logic such that simulation of (1) misalignment, (2) missing section, (3) misalignment + missing section, (4) lost section, and (5) lost + missing section are all mutually exclusive.
- **Slip interpolation** (Lee et al. 2021)

- The dominant type of misalignment errors in the current dataset are so-called “slip” misalignments, in which sections at z-1 and z+1 are well-aligned to each other whereas the section at z is misaligned with respect to both the sections above and below. Such slip misalignments mostly arise locally near folds due to imperfect handling in the alignment process.
- To address this, we replaced the slip-type misalignment simulation of (Lee et al. 2017) by a novel “slip interpolation” (Lee et al. 2021). Specifically, we simulated the slip-type misalignment only in the input, not in the target. The mismatch between the input and target forces the nets to ignore any slip misalignment in the input and produce smoothly interpolated prediction.

### Data and code availability

EM image data and cellular segmentation can be viewed at https://www.microns-explorer.org/cortical-mm3/.

The software tools used to stitch and rough align the dataset is available in the AllenInstitute github repository https://github.com/AllenInstitute/render-modules. The volume assembly process is entirely based on image metadata and transformations manipulations and is supported by the Render service (https://github.com/saalfeldlab/render).

The code for defect detection and resin detection are available at https://github.com/jabae/detectEM/ and the code for running large-scale inference can be found at https://github.com/seung-lab/SEAMLeSS/ under the “alex-emdetector” branch.

The code for alignment using convolutional nets is at https://github.com/seung-lab/SEAMLeSS.

The code for neuronal boundary detection is at https://github.com/seung-lab/DeepEM.

Distributed inference code can be found at https://github.com/seung-lab/chunkflow.

The distributed clustering algorithms for watershed and agglomeration are implemented in https://github.com/seung-lab/abiss, and the workflow management system based on Apache Airflow can be found at https://github.com/seung-lab/seuron.

The code for synaptic partner assignment and synapse segmentation is available at https://github.com/seung-lab/Synaptor/.

CloudVolume, Igneous, and Kimimaro are available at https://github.com/seung-lab/cloud-volume, https://github.com/seung-lab/igneous, and https://github.com/seung-lab/kimimaro.

## Acknowledgements

The authors thank David Markowitz, the IARPA MICrONS Program Manager, who coordinated this work during all three phases of the MICrONS program. We thank IARPA program managers Jacob Vogelstein and David Markowitz for conceiving of and advocating for the MICrONS program. We thank Wade Leonard and Jennifer Wang, IARPA SETAs for their assistance.

The work was supported by the Intelligence Advanced Research Projects Activity (IARPA) via Department of Interior/ Interior Business Center (DoI/IBC) contract numbers D16PC00003, D16PC00004, and D16PC0005. The U.S. Government is authorized to reproduce and distribute reprints for Governmental purposes notwithstanding any copyright annotation thereon. HSS also acknowledges support from NIH/NINDS U19 NS104648, NIH/NEI R01 EY027036, NIH/NIMH U01 MH114824, NIH/NIMH U01 MH117072 NIH/NINDS R01 NS104926, NIH/NIMH RF1 MH117815, NIH/NIMH RF1 MH123400 and the Mathers Foundation, as well as assistance from Google, Amazon, and Intel. XP acknowledges support from NSF CAREER grant IOS-1552868. XP and AT acknowledge support from NSF NeuroNex grant 1707400. AT also acknowledges support from NIMH and NINDS under Award Number U19MH114830.

We thank Stephan Saalfeld, Khaled Khairy and Eric Trautman for help with the parameters for 2D stitching and rough alignment of the dataset.

We thank Garrett McGrath for computer system administration, and May Husseini and Larry and Janet Jackel for project administration at Princeton University.

We thank Brock Wester, William Gray-Roncal, Sandy Hider, Tim Gion, Daniel Xenes, Jordan Matelsky, Caitlyn Bishop, Derek Pryor, Dean Kleissas, Luis Rodriguez and Miller Wilt from John Hopkins University Applied Physics Lab for providing data assessments on the neural circuit reconstruction and infrastructure through BOSSdb We would like to thank the “Connectomics at Google” team for developing Neuroglancer and computational resource donations, in particular J. Maitin-Shepard for authoring neuroglancer and helping to create the reformatted sharded multi-resolution meshes and imagery files used to display the data.

We would like to thank the Google Cloud team, including Keith Binder, Jesus Trujillo Gomez, and Stephen Fang, that supported us during processing.

We would like to thank Amazon and the AWS Open Science platform for providing computational resources. We’d like to also thank Intel for their assistance.

We thank Rob Young for managing the stitching and alignment pipeline at the Allen Institute for Brain Science (AIBS). We thank John Philips, Sill Coulter and the Program Management team at the AIBS for their guidance for project strategy and operations. We thank Hongkui Zeng, Ed Lein, Christof Koch and Allan Jones for their support and leadership. We thank the Manufacturing and Processing Engineering team at the AIBS for their help in implementing the EM imaging and sectioning pipeline. We thank Brian Youngstrom, Stuart Kendrick and the Allen Institute IT team for support with infrastructure, data management and data transfer. We thank the Facilities, Finance, and Legal teams at the AIBS for their support on the MICrONS contract. We thank the Allen Institute for Brain Science founder, Paul G. Allen, for his vision, encouragement and support.

Disclaimer: The views and conclusions contained herein are those of the authors and should not be interpreted as necessarily representing the official policies or endorsements, either expressed or implied, of IARPA, DoI/IBC, or the U.S. Government.

