## Supplementary figures and images for "Petascale neural circuit reconstruction: automated methods"

### Movie S1

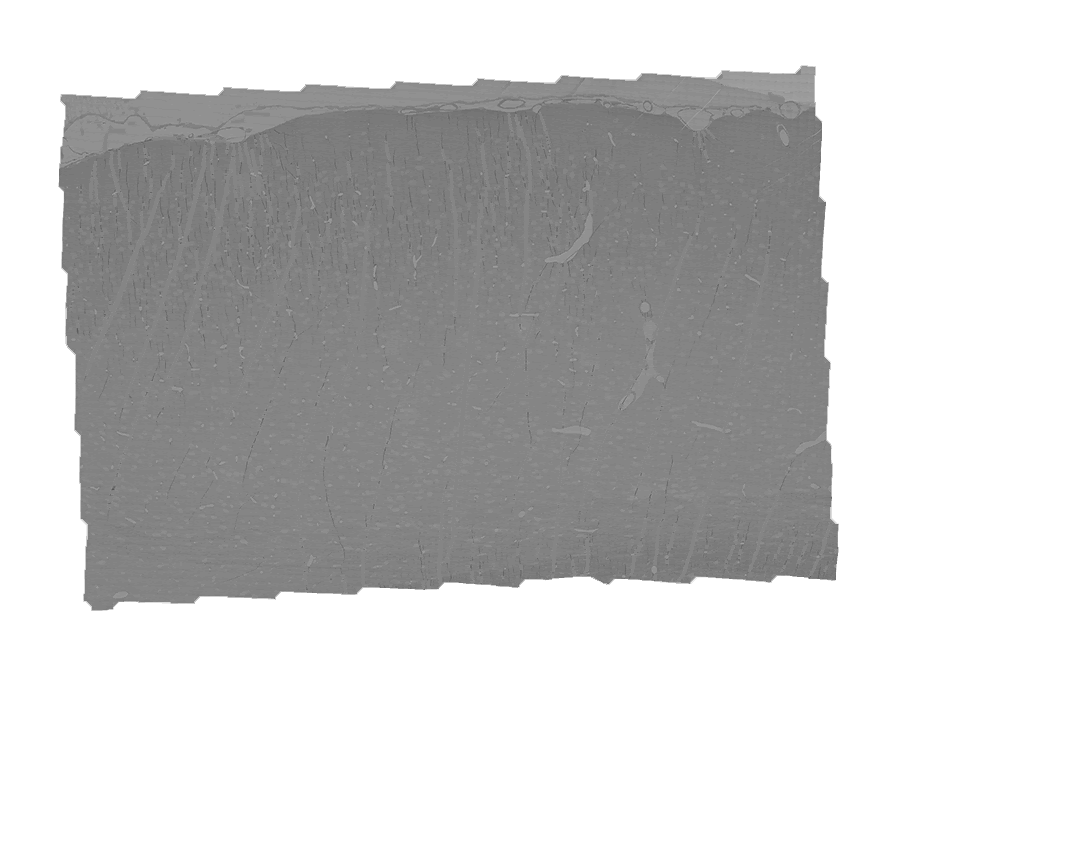
